# A gene-rich mitochondrion with a unique ancestral protein transport system

**DOI:** 10.1101/2024.01.30.577968

**Authors:** David Moreira, Jazmin Blaz, Eunsoo Kim, Laura Eme

## Abstract

Mitochondria originated from an ancient endosymbiotic event involving an alphaproteobacterium^1–3^. Over time, these organelles reduced their gene content massively, with most genes being transferred to the host nucleus before the last eukaryotic common ancestor (LECA)^4^. This process has yielded varying gene compositions in modern mitogenomes, including the complete loss of this organellar genome in some extreme cases^5–14^. At the other end of the spectrum, Jakobids harbor the largest mitogenomes, encoding 60-66 proteins^8^. Here, we introduce the mitogenome of *Mantamonas sphyraenae*, a protist from the deep-branching CRuMs supergroup^15,16^. Remarkably, it boasts the most gene-rich mitogenome outside of jakobids, by housing 91 genes, including 62 protein-coding ones. These include rare homologs of the four subunits of the bacterial-type cytochrome c maturation system I (CcmA, CcmB, CcmC, and CcmF), alongside a unique ribosomal protein S6. During the early evolution of this organelle, gene transfer from the proto-mitochondrial endosymbiont to the nucleus became possible thanks to systems facilitating the transport of proteins synthesized in the host cytoplasm back to the mitochondrion. In addition to the universally found eukaryotic protein import systems, jakobid mitogenomes were reported to uniquely encode the SecY transmembrane protein of the bacterial Type II secretion system; its evolutionary origin was however unclear. The *Mantamonas* mitogenome not only encodes SecY but also SecA, SecE, and SecG, making it the sole eukaryote known to house a complete mitochondrial Sec translocation system. Furthermore, our phylogenetic and comparative genomic analyses provide compelling evidence for the alphaproteobacterial origin of this system, establishing its presence in LECA.

## Results and Discussion

### *Mantamonas sphyraenae* possesses one of the most gene-rich mitogenomes

*Mantamonas sphyraenae* is a protist species belonging to the CRuMs supergroup^15,16^. Our analysis of its complete mitochondrial genome revealed a 53,051 bp long molecule (Figure 1A) with an A+T content of 73.45%. This genome is remarkably compact, with coding regions representing 90.3% of the whole sequence, a value much higher than that of *Diphylleia rotans* (65.9%), the only mitogenome from CRuMs previously characterized^9^, and more similar to those found in most jakobids (89-93%)^8^. We detected many palindromic sequences in intergenic (39) and intragenic (65) regions, as also observed in *D. rotans*, which displays 83 palindromic repeats^9^. However, these A+T-rich (93%) sequences were shorter than in *D. rotans* (8-10 vs 21-45 bp long, respectively) and did not show conserved motifs. The *M. sphyraenae* mitogenome contains no introns, mobile elements, or split genes, and encodes 91 genes, including three rRNAs, 26 tRNAs (which recognize codons corresponding to all the amino acids), the catalytic RNA rnpB of RNase P, and 62 protein-coding genes, as well as one open reading frame (ORF) with no detectable homologs (Figure 1A). The protein-coding gene count overlaps with what is found in jakobids (between 60 and 66), and substantially exceeds that of the recently described gene-rich mitogenomes from *Meteora*^*17*^, nibblerids^18^, and *Microheliella*^*19*^, which contain 50-53 protein-coding genes. *Mantamonas*, therefore, possesses one of the most gene-rich mitogenomes among eukaryotes (Figure 1B).

**Figure 1.**
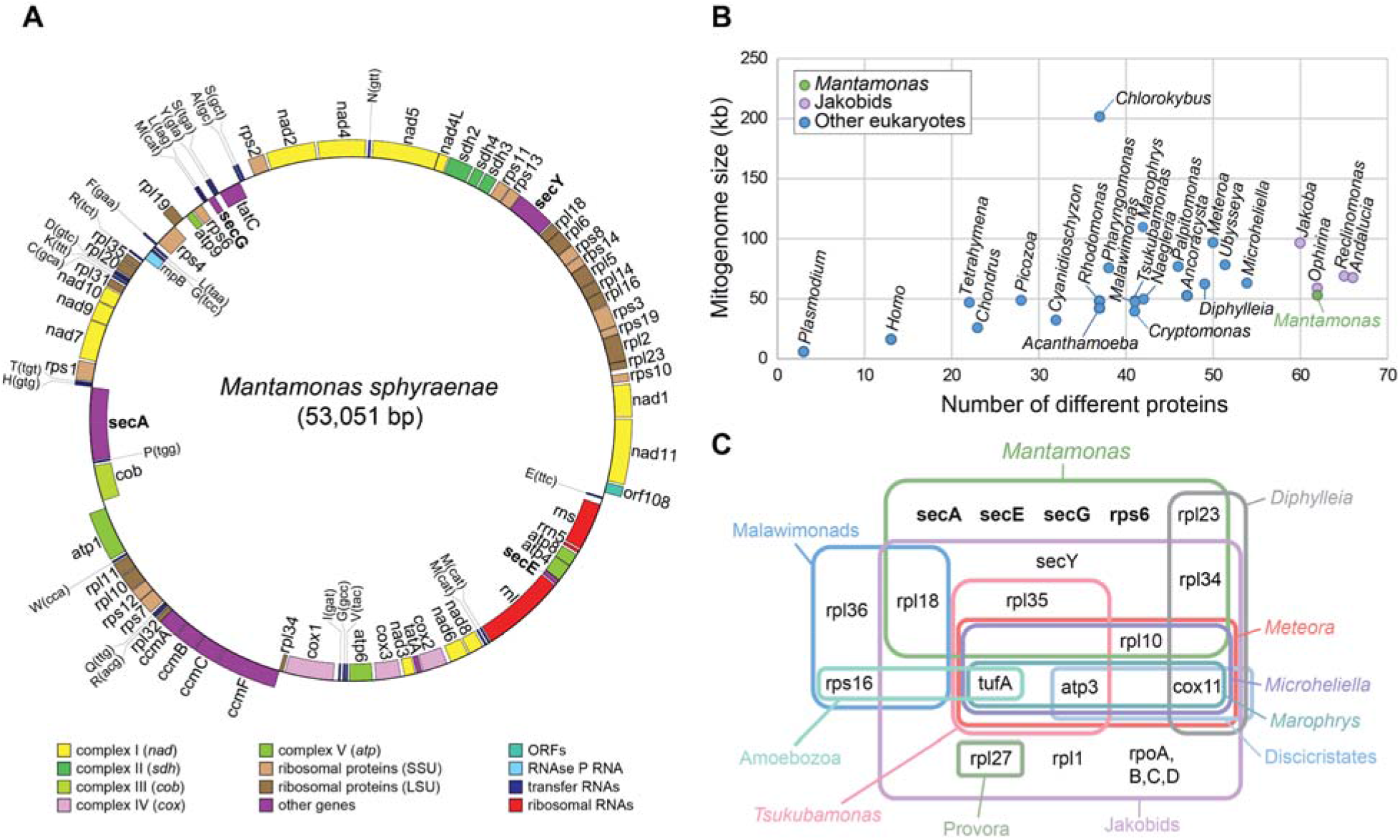
*Mantamonas sphyraenae* possesses one of the most gene-rich mitogenomes. (A) Physical map of the *M. sphyraenae* mitochondrial genome. Genes are color-coded based on their function. Genes on the outer circle are transcribed clockwise; genes on the inner circle counter-clockwise. (B) Size and number of unique protein-coding genes in selected mitogenomes across eukaryotes. (C) Venn diagram of the presence of rare protein-coding genes in selected gene-rich mitochondrial genomes.

With 49 protein-coding genes, *D. rotans* also has a gene-rich mitogenome^9^ compared to most eukaryotes, but significantly below the 62 protein-coding genes found in *Mantamonas*. Interestingly, these two species are the most distant representatives of the CRuMs^15,16^. *Mantamonas sphyraenae* and *D. rotans* are the only eukaryotes that encode the ribosomal protein rpl23 in their mitogenomes, a shared character that provides additional support for the monophyly of CRuMs (Figure 1C). *Diphylleia* also encodes the protein Cox11, which is absent in *Mantamonas*. Apart from that, the mitochondrial gene repertoire of *Diphylleia* is essentially a subset of that of *Mantamonas*. In fact, the *Mantamonas* mitogenome encodes several proteins rarely found in other eukaryotes. These include ribosomal (rpl10, rpl18, and rpl35), cytochrome c maturase (CcmA, CcmB, CcmC, and CcmF), twin-arginine translocase (TatA) proteins, and, remarkably, the SecY translocase, which was previously only found in jakobids^20^ (Figure 1C). The *Mantamonas* mitogenome also codes for the catalytic RNA of the ribonucleoprotein particle that processes tRNA-ends (RNPB)^21^; this mitochondrial gene is only present in a few fungi and protists, including most jakobids^8,22^. Therefore, despite their very large phylogenetic distance, the *Mantamonas* mitogenome exhibits a gene complement very similar to that of jakobids, with the important exception of the absence of the four-subunit bacterial-like RNA polymerase that remains exclusive of jakobids. By contrast, *Mantamonas* possesses the only mitochondrial genome encoding the ribosomal protein S6 (Figures 1C and 2).

Among the rare mitochondrial-encoded proteins found in *Mantamonas*, the four subunits of the bacterial-type cytochrome c maturation system I (CcmA, CcmB, CcmC, and CcmF) have interesting evolutionary implications. They are absent in most eukaryotic lineages, in which they have been replaced by a nucleus-encoded system III (holocytochrome c synthase, HCCS) that does not have prokaryotic homologs^23^. Cavalier-Smith^24^ argued that the ancestral bacterial cytochrome c biogenesis mechanism was inherited from the alphaproteobacterial ancestor of mitochondria in excavates and replaced by the new HCCS in an ancestor of the rest of eukaryotes, supporting the idea that the root of the eukaryotic tree is situated on the excavate branch. However, the presence of the four Ccm subunits in Mantamonas supports that both systems must have coexisted in the branch leading to the Amorphea for a longer time than previously thought. Moreover, the recent discovery that *Ancoracysta* has both a mitochondrion-encoded system I and a nucleus-encoded system III, seemingly acquired by horizontal gene transfer, argues against a single acquisition of HCCS in eukaryotes that could be used to pinpoint the position of the eukaryotic root^12^. The presence reported here of the four Ccm subunits in *Mantamonas* supports that both systems must have coexisted in the branch leading to the Amorphea for a longer time than previously thought.

### A complex ancestral protein transport system in *Mantamonas* mitochondria

Jakobid mitochondria possess a rich repertoire of genes involved in synthesizing protein translocation systems, including the TAT translocase and the SecY transmembrane protein of the bacterial Type II protein secretion system^20^. Whereas the TAT proteins are also encoded in mitogenomes of other eukaryotic groups^3^, SecY has never been found outside the jakobids^8^ (Figure 2). Because the jakobid SecY homologs are encoded on mitogenomes, it can be hypothesized that they were inherited from the alphaproteobacterial ancestor of mitochondria. However, these jakobid SecY sequences are extremely divergent, and previous phylogenetic analyses have failed to show their expected relationship with alphaproteobacterial homologs, making their evolutionary origin unclear^25^. By contrast, including the more conserved *Mantamonas* SecY in our Maximum Likelihood (ML) phylogenetic inference produced a tree that shows the monophyly of all the mitochondrial SecY and strong support for their alphaproteobacterial origin (Figures 3A and S1). Interestingly, a 14-gene genomic region in which the gene *secY* is nested shows perfect synteny conservation between the mitogenomes of *Mantamonas* and the jakobid *Reclinomonas americana* (Figure 3B). This provides strong additional support for the common origin of this gene in both lineages in the mitochondrion of an ancestor of the two groups.

**Figure 2.**
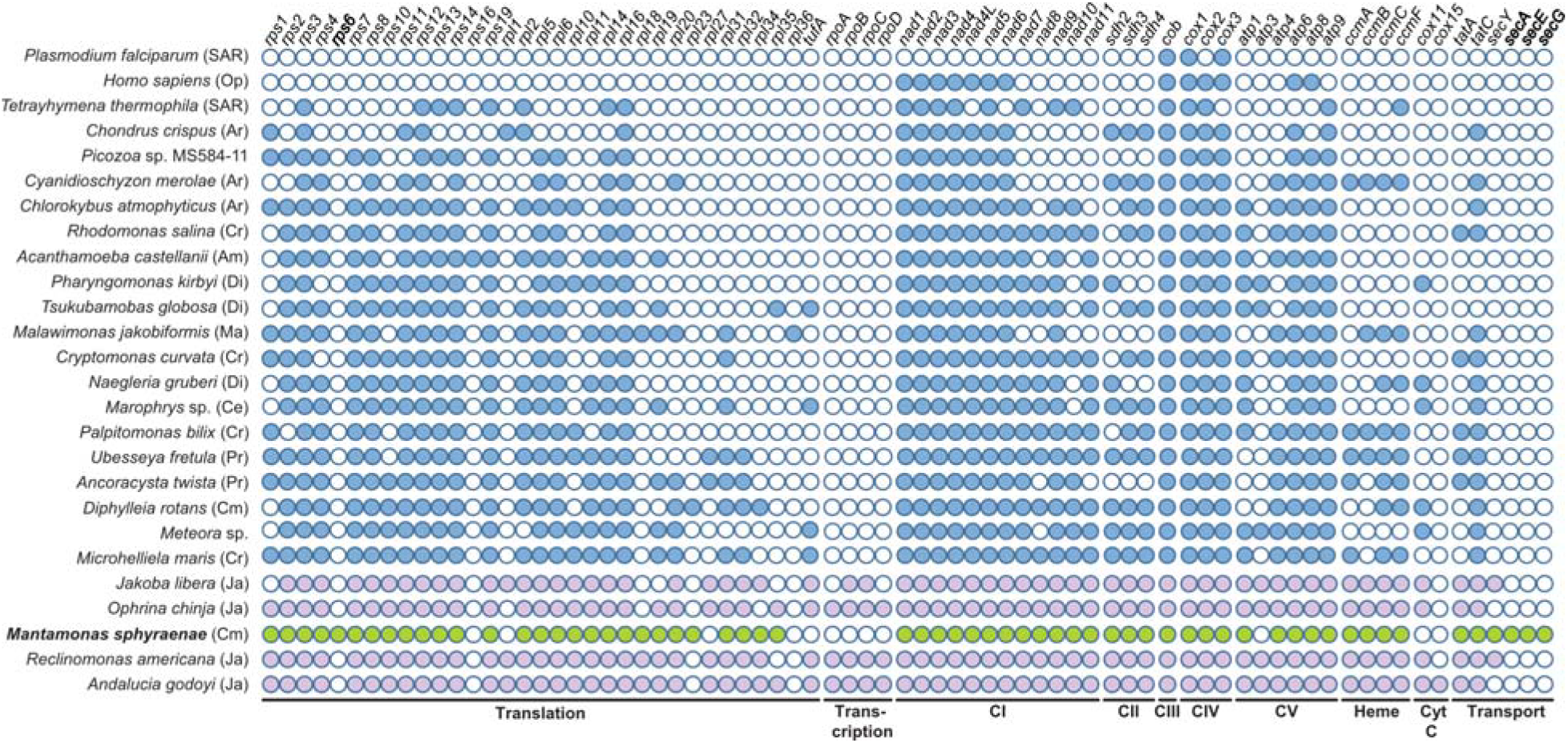
Protein-coding capacity of mitochondrial genomes across eukaryotes. Gene presence is indicated by a filled circle and is derived from ^17^ updated with additional lineages. The species are arranged in descending order based on the number of protein-coding genes in their mitochondrial genome. Jakobids are depicted in pink, *Mantamonas* in green and other eukaryotes in blue. Genes in bold have been uniquely found in the *Mantamonas* mitogenome. SAR: Stramenopiles, Alveolata, Rhizaria; Ce: Centrohelids; Cr: Pancryptista; Ar: Archaeplastida; Pr: Provora; Cm: CRuMs; Op: Opisthokonta; Am: Amoebozoa, Ma: Malawimonadida, Di: Discoba. CI-CV - electron transport chain complex I–V. Cyt C: Cytochrome c oxidase.

**Figure 3.**
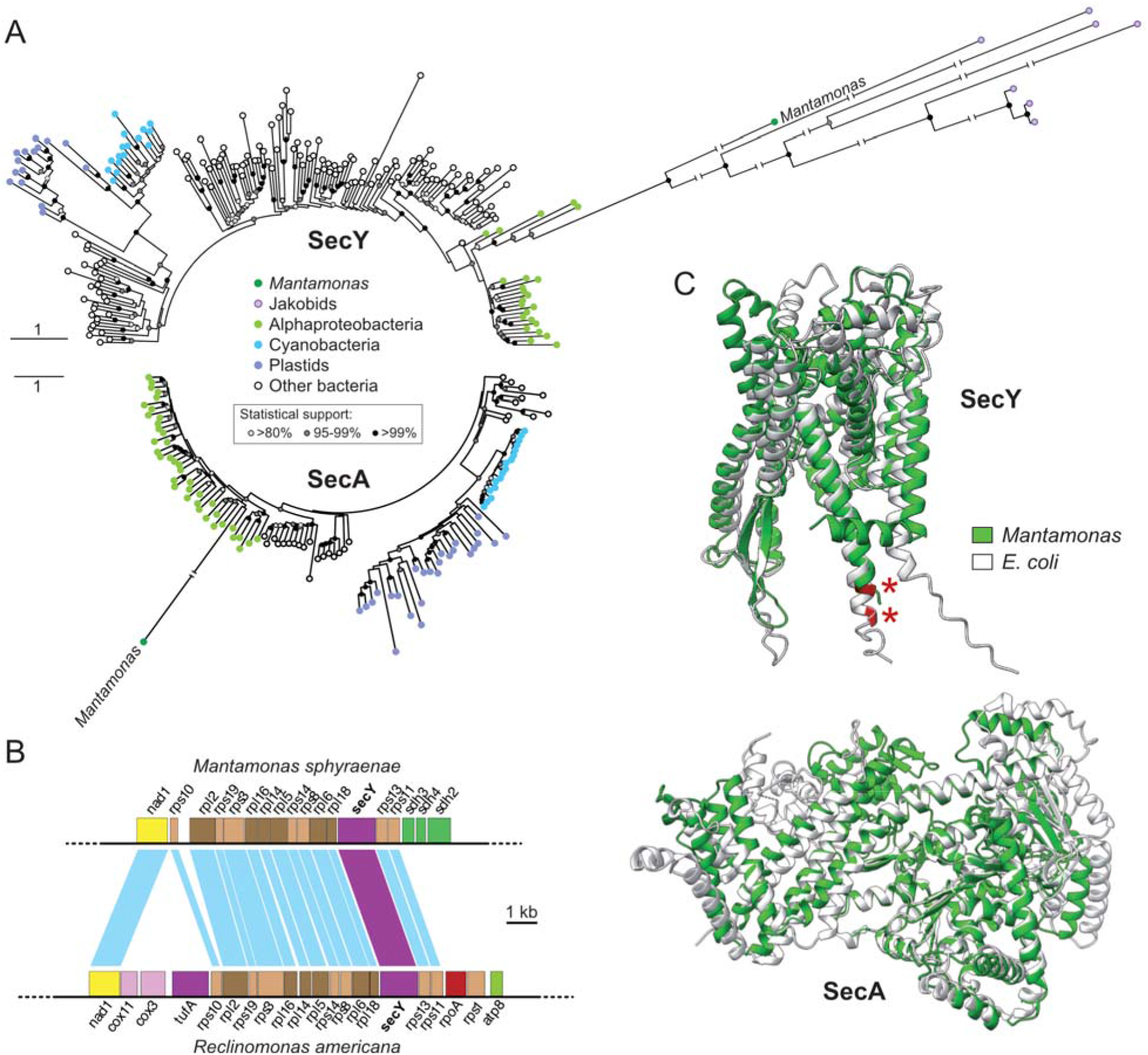
An ancestral bacterial Sec system in Mantamonas. (A) Maximum likelihood phylogenetic trees of SecY (183 sequences, 406 sites) and SecA (129 sequences, 920 sites). (B) Conservation of synteny around the *secY* gene in the mitochondrial genomes of *M. sphyraenae* and *R. americana*. Homologous genes are connected by blue lines, except the *secY* gene, highlighted in purple. (C) Predicted structures of *Mantamonas* SecY and SecA proteins (green) superposed to their *E. coli* homologs (white). The C-terminal KK dibasic motif in both SecY structures is indicated by red asterisks.

In bacteria, the Sec system is the principal protein translocase. For example, it translocates 96% of the exported polypeptides in *Escherichia coli*^*26*^. SecY is accompanied by several other proteins to form a complete active Sec translocation system. They include the chaperone SecB, the ATPase SecA, and the membrane proteins SecD, SecE, SecF, and SecG^27^. However, these other Sec proteins have not been identified in the mitochondrial or nuclear genome sequences of jakobids, so the function of their SecY protein remains elusive^25,28^. By contrast, a major difference between the mitogenomes of jakobids and *Mantamonas* is the exceptional presence of genes encoding SecA, SecE and SecG in the latter (Figure 1). SecA uses energy from ATP hydrolysis to move hydrophilic proteins through the membrane by repeated cycles of SecA membrane insertion and deinsertion^29^, whereas SecY, SecE, and SecG form a membrane-embedded channel, the SecYEG complex^30^. These four proteins have been shown to be enough to build a Sec system with strong protein translocation activity in *E. coli*^*30,31*^. *Mantamonas* is the only eukaryote where mitochondrial copies of the genes encoding all these proteins have been identified so far (Figure 2). This system is likely functional despite the absence of SecD and SecF. Indeed, these two proteins are not essential for the Sec activity in *E. coli* and many bacteria lack the genes encoding them^32^.

Like for SecY, our ML phylogenetic analyses strongly support the alphaproteobacterial origin of *Mantamonas* SecA (Figures 3A and S2), which has a well-conserved structure when compared to its bacterial homologs (Figure 3C). By contrast, SecE and SecG are much smaller proteins, and the sequence similarity of the *Mantamonas* sequences to their prokaryotic homologs was too low to yield resolved phylogenies. Nevertheless, their presence in the *Mantamonas* mitogenome combined with the clear alphaproteobacterial origin of the genes encoding SecY and SecA supports the hypothesis that *Mantamonas* has kept a functional SecYEG-SecA system inherited from the protomitochondrial endosymbiont.

We hypothesize that the soluble motor protein SecA drives the passage of the polypeptide through the channel made by the integral membrane protein SecY. Significantly, the SecY protein of *Mantamonas* has the C-terminal dibasic motif KK (Figure 3C) involved in translocation coupling with SecA^33,34^, which has been lost in most jakobid SecY sequences. We looked for possible candidate proteins to be translocated by the *Mantamonas* SecYEG-SecA complex. As mentioned above, it has been proposed that Cox11 is the target of SecY in jakobids^20^, but this protein is not encoded in the *Mantamonas* mitogenome. Nevertheless, jakobids have other mitochondrial proteins with predicted bacterial-type signal peptides that may be alternative candidates (e.g., nine proteins in *R. americana*^*20*^). Surprisingly, in *Mantamonas* only two mitochondrial-encoded proteins, Cox2 and Atp4, are predicted to have a signal peptide (Figures 4A and S3), being therefore the most likely candidates to be the substrate of the SecYEG-SecA complex. It has been proposed that the Sec system is especially efficient for the translocation of proteins with large hydrophilic domains^35^. In agreement, among the *Mantamonas* mitochondrial membrane proteins, Cox2 and Atp4 have the longest hydrophilic domains (Figure 4B). Consistently, these two proteins, which are very frequently encoded in mitogenomes of various eukaryotes (Figure 2), do not contain a signal peptide in the SecY-lacking *D. rotans*.

**Figure 4.**
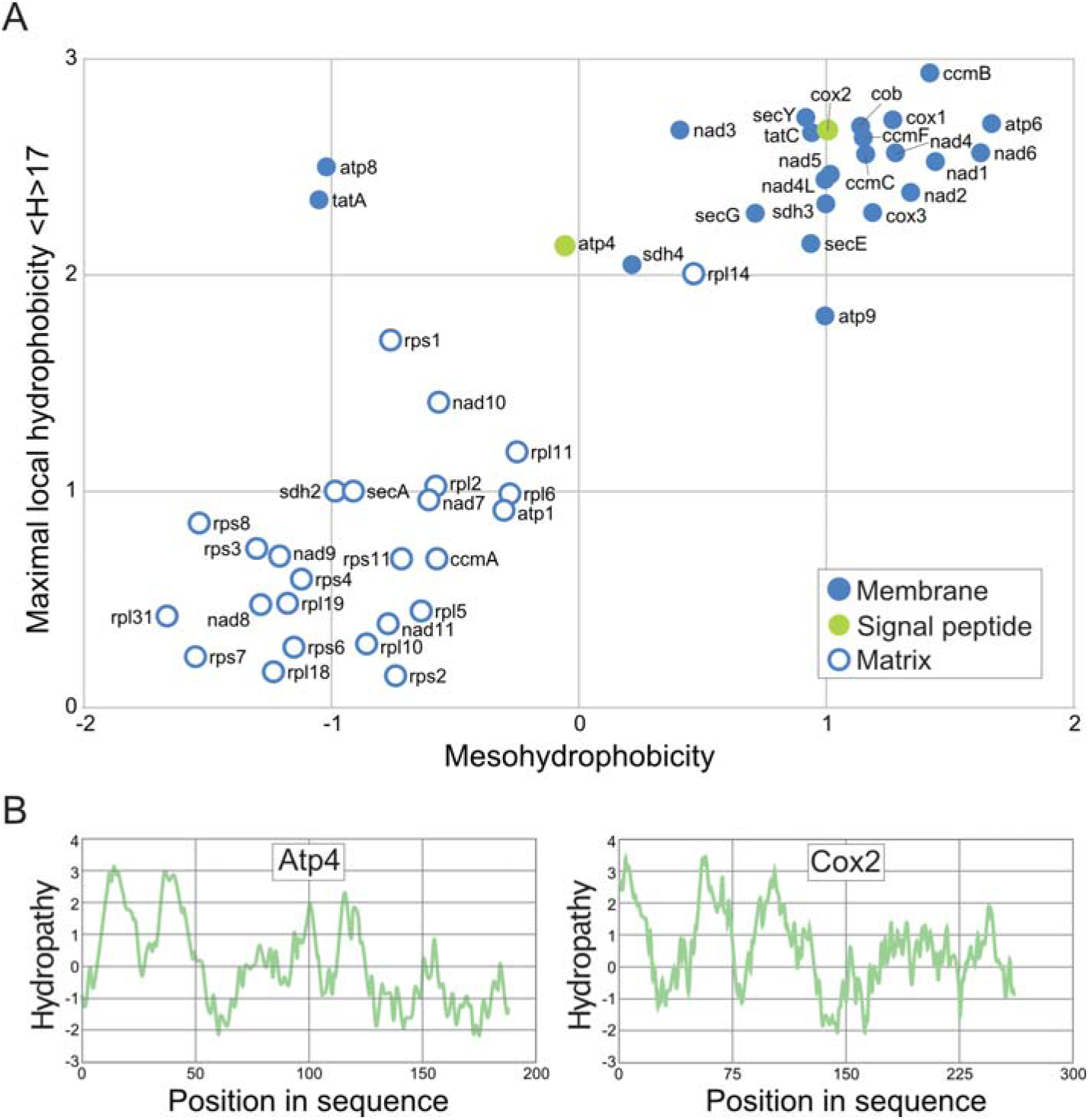
Hydrophobicity of mitochondrial proteins of Mantamonas. (A) Hydropathy analysis of the predicted protein sequences encoded in the mitogenome of *M. sphyraenae*. Proteins are color-coded according to their predicted location and the presence of a signal peptide. (B) Hydropathy plots of the proteins Atp4 and Cox2, potential substrates of the *M. sphyraenae* mitochondrial Sec system.

## Convergent gene losses dominated post-LECA mitochondrial evolution

Even the gene-richest mitogenomes, such as those of jakobids and *Mantamonas*, have a dramatically smaller gene content than free-living or even parasitic alphaproteobacteria. Comparative genomics has shown that this difference can be explained by a combination of gene loss and gene transfer to the host nucleus4. As a result, contemporary mitogenomes retain a relatively small number of genes, most of which can be classified into two main categories: genes involved in protein synthesis (mostly rRNA, tRNA, and ribosomal protein genes) and genes involved in the electron transport chain (many of them coding for membrane proteins). It is assumed that a large mitogenome reduction occurred before LECA12,36,37, although in some lineages, there is clear evidence for more recent reduction episodes that led to tiny mitogenomes, those found in apicomplexan parasites being a dramatic example38, or even to complete loss of mtDNA in a variety of anaerobic protists11,13. However, the tempo of mitochondrial genome reduction that occurred between LECA and the diversification of the major eukaryotic phyla remains uncertain. The discovery of gene-rich mitogenomes in jakobids led to the hypothesis that they were early-diverging eukaryotes emerging near the root of the eukaryote tree39,40. According to this idea, all other eukaryotes would have experienced a genome reduction after the divergence of jakobids and before their diversification into the rest of contemporary lineages. However, the position of the root of the eukaryote tree remains debated41–43, and the discovery of other distantly related, non-jakobid gene-rich mitogenomes9,12,44, now further complemented with the Mantamonas one, suggests that the evolution of gene content in mitochondrial genomes has been more complex than previously hypothesized and involved a large number of convergent gene losses. As a consequence, a rich gene content is not enough evidence to make hypotheses regarding the position of the root or the deep relationships among eukaryotic groups.

Another discovery with important evolutionary implications is the presence of the SecYEG-SecA system in Mantamonas. SecY was already known in jakobids (Tong et al. 2011), but its origin remained unclear because the jakobid sequences appeared not to be related to their alphaproteobacterial homologs, opening the possibility for a late acquisition by HGT from an unknown bacterial donor (Petrů et al. 2021). In the case of Mantamonas, both SecY and SecA have a clear alphaproteobacterial origin (Figure 3A). Furthermore, the addition of the slower-evolving Mantamonas SecY combined with complex evolutionary models helped resolve the phylogeny and strongly suggests that the jakobids SecY homologs have a common origin with the Mantamonas one, and descend from their alphaproteobacterial counterpart. Given the large phylogenetic distance between jakobids and CRuMs, this result supports that this protein translocation system was present in the mitochondrion of LECA and kept during the early diversification of contemporary eukaryotic lineages, to be subsequently lost in most of these lineages several times independently.

The presence of several mitochondrial protein translocases in LECA (including TIM/TOM, Oxa, Tat, SAM, miT2SS, and Sec) likely introduced some functional redundancy and potentially relaxed the selective pressure on these systems. This not only accounts for the current mosaic distribution of some of them but also certain lineage-specific dissimilarities observed in the universal TIM/TOM complex (Lister et al. 2005). The evolution of these systems provides a striking example of the greater complexity of the LECA mitogenome compared to contemporary eukaryotes, which have undergone varying degrees of reduction over time. Sequencing mitogenomes of lesser-known eukaryotic lineages is expected to contribute more examples, further elucidating the complexity of the ancestral mitogenomes.

## Methods

### Assembly of *Mantamonas sphyraenae* mitochondrial DNA

The genome sequence of *M. sphyraenae* was generated as described in Blaz *et al*.^*16*^. Briefly, whole DNA from this protist was sequenced using both long (PacBio) and short (Illumina) read sequencing, with a coverage of 112x for PacBio and 115x for Illumina. A hybrid assembly was generated using FALCON^45^ and the mitogenome was identified in this assembly by looking for contigs containing typical mitochondrial genes (e.g., *cox* and *nad* genes) using BLAST^46^. All these genes were found in a single 53,051 bp long contig.

### Mitogenome gene prediction and annotation

Mitochondrial gene prediction and annotation was done using MFannot v1.36 (https://megasun.bch.umontreal.ca/apps/mfannot/)^47^ with the standard genetic code. All annotations and gene boundaries were inspected manually. Several small divergent ORFs (e.g., the gene coding for SecG) were annotated using PROST^48^ and InterproScan v5.31-70.0^49^. The graphical representation of the annotated mitogenome was generated using OrganellarGenomeDRAW (OGDRAW) v1.3.1^50^. Palindromic sequence elements were detected by EMBOSS explorer (http://emboss.bioinformatics.nl/cgi-bin/emboss/palindrome) with the following parameters: Minimum and maximum length of palindromes was 6 and 100, respectively; the maximum gap between elements was 10, and no mismatch was allowed in palindrome.

### Hydrophobicity analysis and protein structure predictions

Signal peptides, transmembrane helices, and protein localization were predicted using TOPCONS v2.0 ^51^ and DeepTMHMM v1.0.24^52^. Graphical representation of proteins was done using Protter^53^. Individual protein hydropathy plots were generated with ProtScale (https://web.expasy.org/protscale/) applying the Kyte and Doolittle method^54^ with a window size of 9. Mesohydrophobicity versus maximal local hydrophobicity (<H>17) values were calculated for the whole set of mitochondrion-encoded proteins using MitoProt II^55^.

Protein structures were predicted using AlphaFold2 with default parameters^56^. Predicted *Escherichia coli* protein structures were downloaded from the AlphaFold DB^57^. Alignment of protein structures was done with ChimeraX^58^.

### Phylogenetic analyses

Protein sequences similar to the *Mantamonas* mitochondrial SecY and SecA were retrieved using BLAST^46^ searches against the RefSeq^59^ and GTDB^60^ databases. Sequences were aligned with MAFFT L-INS-i v7.450^61^ and trimmed using trimAl^62^ and TCS^63^. Maximum likelihood phylogenetic trees were then reconstructed using the LG+C60+F+Γ4 model of sequence evolution in IQ-TREE^64^. Statistical support was obtained with 1000 ultrafast bootstraps^65^.

## Data and code availability

The read data associated with the mitochondrial genome of *Mantamonas sphyraenae* have been submitted to the NCBI SRA database (Bioproject number PRJNA886733). The prediction of protein-coding genes and RNAs is available at Figshare (10.6084/m9.figshare.24635520).

## Acknowledgments

This project has received funding from the European Research Council (ERC) under the European Union’s Horizon 2020 research and innovation programme (ERC Starting Grant No 803151 to L.E. and ERC Advanced Grant No 787904 to D.M.) and the Simons Foundation (Grant No 876199 to E.K.).

## Author contributions

Conceptualization, D.M., and L.E.; resources, E.K.; formal analysis, D.M, J.B., E.K, and L.E; writing – original draft, D.M., and L.E..; writing – review & editing, all; supervision, D.M., E.K. and L.E.; funding acquisition, D.M., E.K, and L.E.

## Supplemental Information

**Supplementary Figure S1. Complete SecY phylogenetic tree**. Maximum likelihood phylogenetic tree based on SecY sequences (183 taxa, 406 sites). Statistical support indicated on the branches corresponds to 1,000 ultra-fast bootstrap replicates. The scale bar indicates the number of substitutions per site.

**Supplementary Figure S2. Complete SecA phylogenetic tree**. Maximum likelihood phylogenetic tree based on SecA sequences (129 taxa, 920 sites). Statistical support indicated on the branches corresponds to 1,000 ultra-fast bootstrap replicates. The scale bar indicates the number of substitutions per site.

**Supplementary Figure S3. Graphical representation of Cox2 and Atp4 predicted signal peptides and insertion into the inner mitochondrial membrane**. IMS: intermembrane mitochondrial space; TM: transmembrane. Generated by Protter and modified.

