## Supplementary Figure S1 for "A gene-rich mitochondrion with a unique ancestral protein transport system"

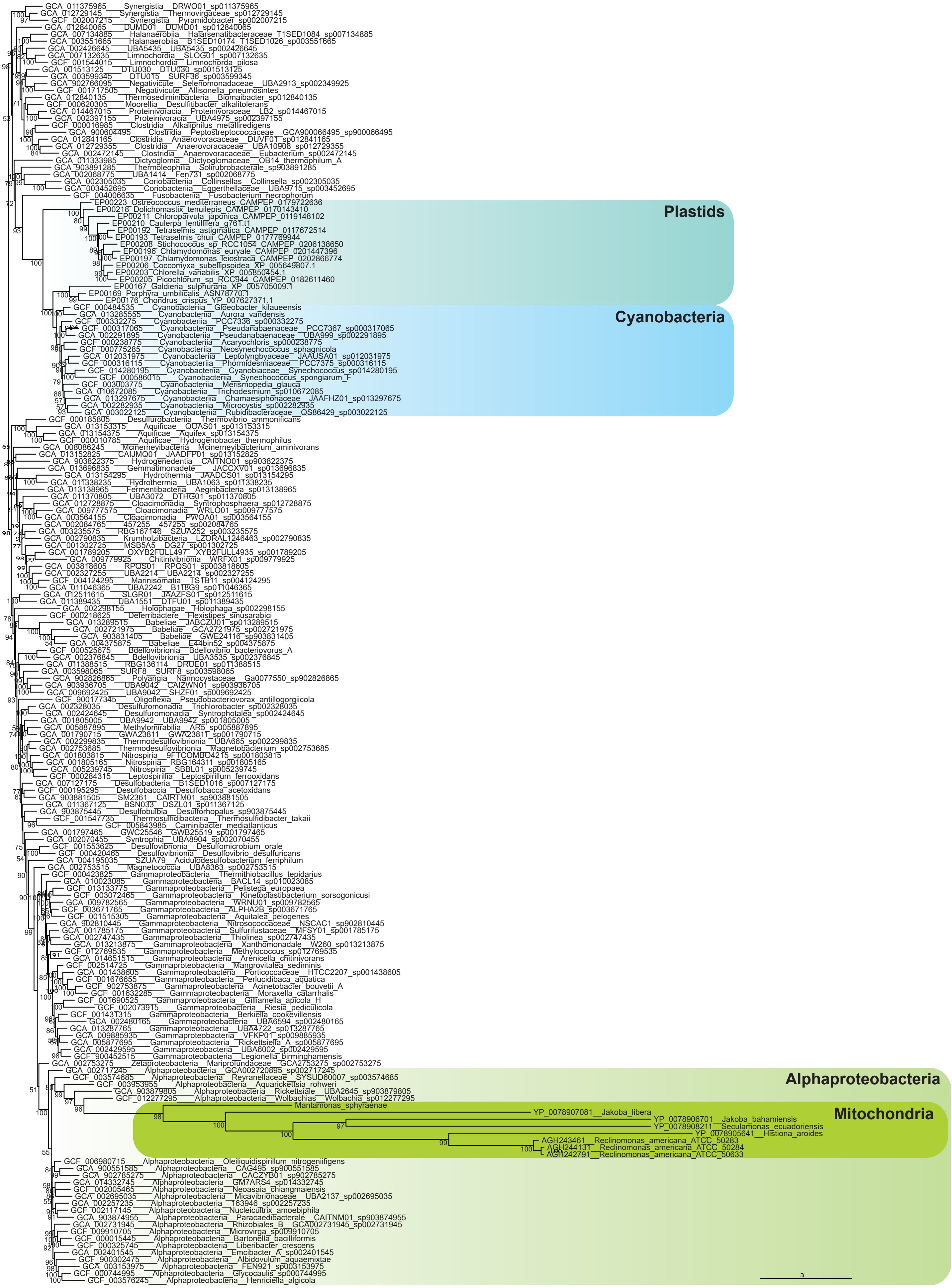


**Figure S1. Complete SecY phylogenetic tree.** Maximum likelihood phylogenetic tree based on SecY sequences (183 taxa, 406 sites). Statistical support indicated on the branches corresponds to 1,000 ultra-fast bootstrap replicates. The scale bar indicates the number of substitutions per site.
