## Supplementary Figure S2 for "A gene-rich mitochondrion with a unique ancestral protein transport system"

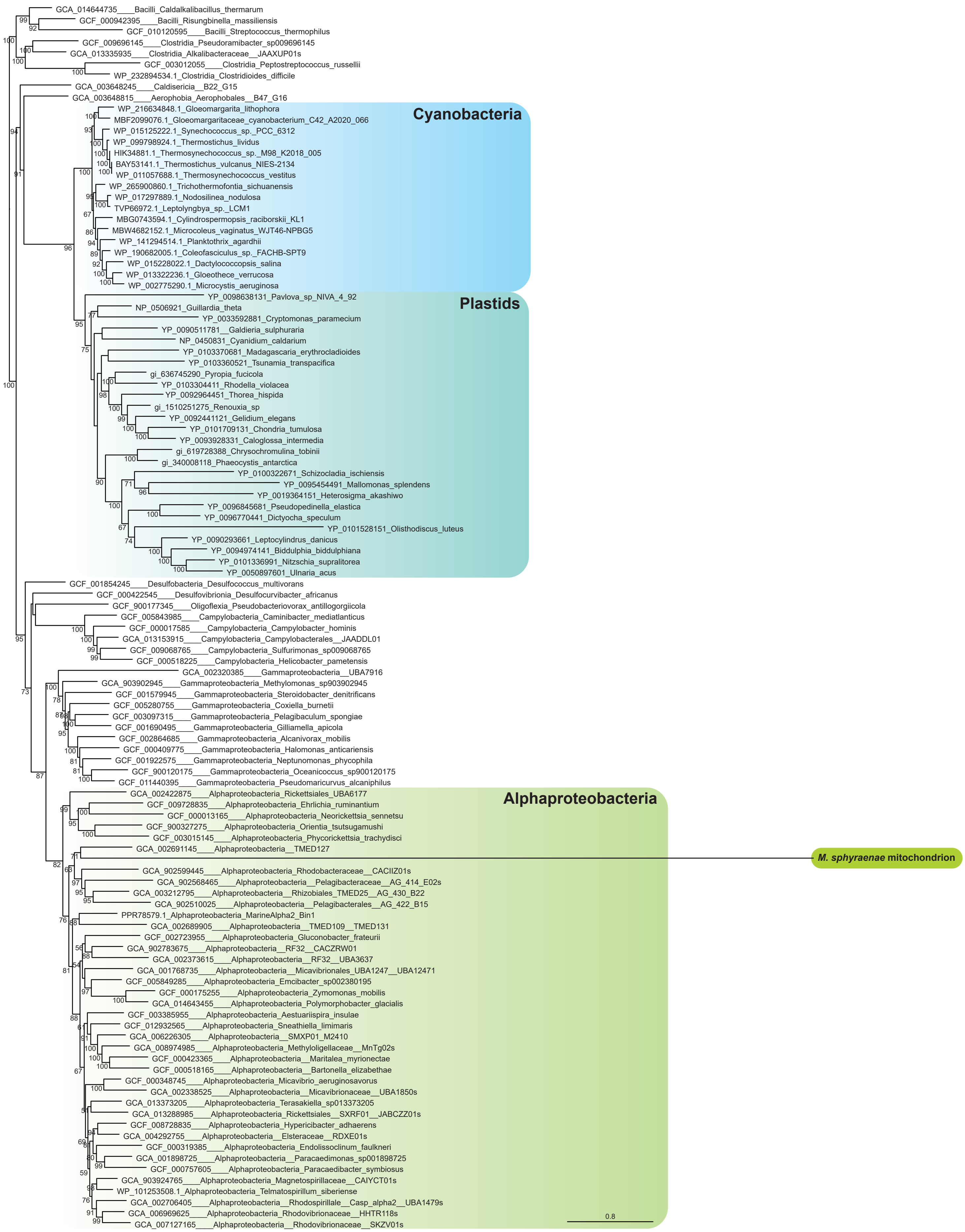


**Figure S2. Complete SecA phylogenetic tree.** Maximum likelihood phylogenetic tree based on SecA sequences (129 taxa, 920 sites). Statistical support indicated on the branches corresponds to 1,000 ultra-fast bootstrap replicates. The scale bar indicates the number of substitutions per site.
