## Supplementary Figure S3 for "A gene-rich mitochondrion with a unique ancestral protein transport system"

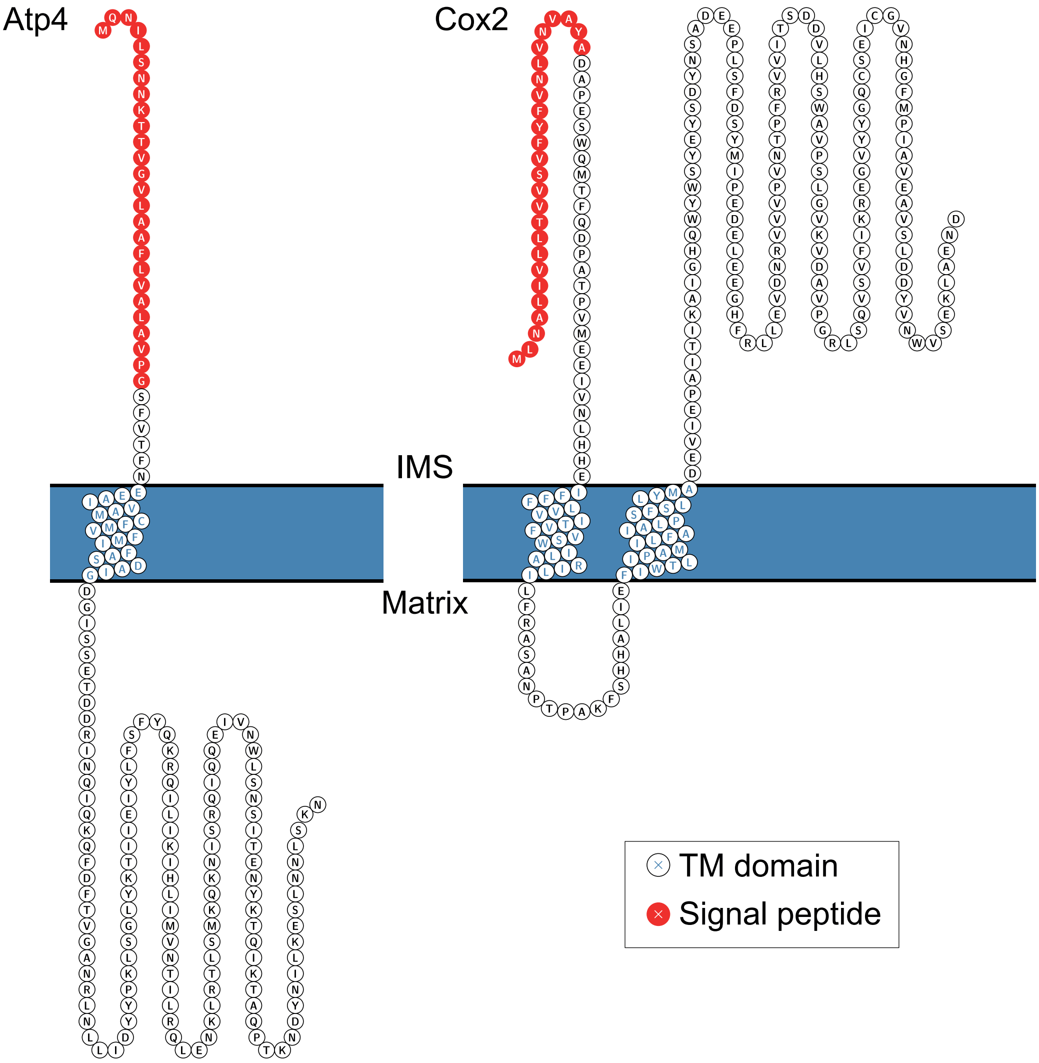


**Figure S3. Graphical representation of Cox2 and Atp4 predicted signal peptides and insertion into the inner mitochondrial membrane.** IMS: intermembrane mitochondrial space; TM: transmembrane.
